# MuStARD: a Deep Learning method for intra- and inter- species scanning identification of small RNA molecules

**DOI:** 10.1101/547679

**Authors:** Georgios K Georgakilas, Andrea Grioni, Konstantinos G Liakos, Eliska Malanikova, Fotis C Plessas, Panagiotis Alexiou

## Abstract

Genomic regions that encode small RNA genes exhibit characteristic patterns in their sequence, secondary structure, and evolutionary conservation. Deep Learning algorithms are efficient at classifying examples based on such learned patterns. Here we present MuStARD (gitlab.com/RBP_Bioinformatics/mustard) a Deep Learning framework that can learn patterns associated with user-defined sets of genomic regions, and scan large genomic areas for novel regions exhibiting similar characteristics. We demonstrate that MuStARD can be trained on different classes of human small RNA loci (pre-miRNAs and snoRNAs) and outperform state of the art methods specifically designed for each specific class. Furthermore, we demonstrate the ability of MuStARD for inter-species identification of functional elements by predicting mouse small RNAs (pre-miRNAs and snoRNAs) using models trained on the human genome. MuStARD is easy to deploy and extend to a variety of genomic classification questions.

## INTRODUCTION

Since the human genome was first sequenced about two decades ago (Lander et al. 2001), our understanding of regulatory and non-coding elements in the human and other model organisms has been steadily increasing (ENCODE Project Consortium 2012) with the identification and cataloguing of a variety of coded molecule and regulatory region classes. Non-coding RNA Molecule families such as microRNA (miRNA), Small nucleolar RNA (snoRNA), Small nuclear RNA (snRNA), Piwi-interacting RNA (piRNA), Short hairpin RNA (shRNA), Small interfering RNA (siRNA), Long non-coding RNAs (lncRNA) and others, now populate the functional expression map of known genomes. Along with our deeper understanding of well-established organism genomes, the total number of sequenced genomes has been increasing hand in hand with fast pace. NCBI currently lists just over 7000 eukaryotic sequenced genomes, of which almost 50 have fully assembled genomes, and approximately 1000 have some assembled chromosomes.

The majority of these newly sequenced genomes cannot be experimentally annotated to the depth of gold-standard genomes such as human, mouse, or Drosophila. Computational methods are continuously being developed with the goal of predicting the location of small non-coding RNA loci in genomes. In silico methods utilizing sequence homology are often employed for the annotation of novel genomes, projecting functional regions of well-annotated species to homologous genomic regions of less annotated genomes. For example, covariant model approaches (Nawrocki 2014) identified 529 specific and highly-conserved small RNA families in small-scale genomes (Kalvari et al. 2018). These 529 miRNA family models represent an unknown portion of the 38,589 miRNA sequences (1,917 human) catalogued in miRBase, since each model can be comprised of molecules derived from several species. Despite the usefulness of evolutionary conservation as a feature, pure homology-based approaches will be, by definition, unable to annotate RNA families that do not have a close homologue in model species. For example, many human miRNAs have developed only recently in evolutionary history and can be found only in other primates (estimated 40% (Wang et al. 2011)). Approaches purely based on homology would strongly limit the identification of small RNA for a newly sequenced species with no well-annotated close ‘relatives’.

Beyond homology-based approaches, genomes can be ‘scanned’ for regions of known characteristics, such as a specific motif, or sequence, and their putative function annotated. For these approaches tools specifically trained on a class of small RNA molecules can be used. However, even for well-studied classes such as miRNAs, the accuracy of such programs makes scanning of large genomic regions unrealistic. Over 30 programs aiming at pre-microRNA identification have been developed, but none achieving accurate genome-wide prediction (Saçar Demirci, Baumbach, and Allmer 2017). A large drawback of such methods is their dependence on expert-defined features and background sets. This process involves an arbitrary number of features that have been conceptualized on ad hoc bases, usually derived on empirical data that are interpreted based on personal experiences and assumptions. This ad hoc process of feature extraction frequently introduces biases that might severely affect building robust models while at the same time does not offer the possibility to utilize and also unveil all underlying patterns.

Here, we present a Machine Learning (ML) method that improves the accuracy of non-coding RNA prediction in known species, and demonstrate that the model trained on a user selceted species can be used to scan large genomic regions and identify cross-species functional elements of the same class. We have chosen to apply our method on two different classes of small RNAs: miRNA precursors (pre-miRNAs) and small nucleolar RNAs (snoRNAs). Precursor miRNAs are intermediate RNA molecules of miRNA biogenesis that form stable hairpin structures of approximately 60-100 nucleotides. The first novel miRNAs were identified by sequencing total RNA of their approximate length (Lagos-Quintana et al. 2003; Lee and Ambros 2001; Lau et al. 2001). Based on the characteristics of the first sequenced miRNAs, computational methods were introduced to accelerate the identification process. Current computational methods utilize some combination of manually produced features based on genomic sequence and conservation, as well as predictions of RNA folding. These features could include the free energy of folding, folding stem length, nucleotide content in the stem, occurrence of matching pairs and so on. In a recent thorough comparison of several highly cited programs, it was observed that no tool significantly outperforms all other tools on all tested data sets (Saçar Demirci, Baumbach, and Allmer 2017). Additionally, none of the current tools can employ a ‘scanning’ mode for large genomic regions leading to accurate pre-miRNA loci identification. Currently, the latest miRBase release (Kozomara, Birgaoanu, and Griffiths-Jones 2018), the main repository of known miRNA sequences gives access to 38,589 pre-miRNAs from 271 organisms with 1,917 being of human origin.

The highly competitive field of pre-miRNA prediction can be juxtaposed with the relative scarcity of snoRNA prediction algorithms. Discovered shortly after the sequencing of the human genome (Kiss 2002) snoRNAs play an important role in the processing and modification, of other classes of RNAs. Over ten years ago, the human genome was scanned for snoRNAs using snoSeeker (Yang et al. 2006, 2010) that appears to not be available anymore, identifying approximately 300 snoRNA loci. Hundred more snoRNAs were later identified by small RNA sequencing of diverse species and filtering through a computational algorithm (Yang et al. 2010). However, despite these initial attempts, high-throughput experimental methods have reached the conclusion that the human genome contains a large number of snoRNAs expressed in low levels (Kishore et al. 2013). For the well-annotated human genome, the use of large experimental data from ENCODE allowed the further identification of snoRNAs, to almost double the number of known loci to ~1000 (Jorjani et al. 2016). Identification of snoRNAs in other species has proceeded in a slow pace and with severe setbacks (Makarova and Kramerov 2011) as resources previously available become obsolete and eventually seize to exist (Lestrade 2006; Xie et al. 2007). The field of snoRNA prediction appears too small to warrant attention of large initiatives to implement complex machine learning architectures and manually curated features. Here, we will demonstrate that MuStARD can accurately predict snoRNA locations, proving that it will be a useful tool for the generalized identification and annotation of less studied classes of snoRNAs. As a demonstration of the predictive power and limitations of our method, we will demonstrate predictions based on training in the entire known snoRNA set, and also predictions based on training in specific families of snoRNAs that exhibit more consistent secondary structures.

Machine Learning describes the field of computer science that involves the development of mathematical models and their implementations with the purpose of enabling computers to learn concepts and patterns embedded in data. Neural Networks are a subfield of ML algorithms with a rich history starting decades ago (Fitch 1944) with attempts to approximate the process of learning in the brain by stacking interconnecting layers of artificial neurons. Deep Learning (DL) is a term that refers to recent breakthroughs in the field of Neural Networks including a collection of new methodologies that have outperformed well-established ML algorithms in various learning tasks (LeCun, Bengio, and Hinton 2015). Deep Neural Networks offer significant flexibility and remarkable accuracy provided enough data, especially for complex learning tasks. A subset of Deep Neural Networks termed Convolutional Neural Networks (CNNs) are able to operate directly on raw data such as DNA/RNA sequences without the need of pre-processing and feature extraction. CNNs use convolutional layers to process the input prior to propagating the signal to the dense part of the network and in the process extract important features by themselves. Even though the power of Deep Learning only became obvious in the past five years, there are already dozens of published studies that applied a plethora of Deep Learning architectures in the various fields of Biology (Ching et al. 2018). For example, epigenomic data were used to infer gene expression (Singh et al. 2016), and ovarian cancer subtypes were defined from gene and microRNA expression as well as DNA methylation (Liang et al. 2015). DeepBind (Alipanahi et al. 2015) was the first application of CNNs in transcription factor binding recognition tasks. DeepSEA (Zhou and Troyanskaya 2015) and DanQ (Quang and Xie 2016) are also CNN-based frameworks that were trained on a large multi-cell-type compendium of chromatin-profiling data, including DNase I sensitivity, TF and histone-mark ChIP-seq data. Basset (Altman et al. 2016) and DeepEnhancer (Xu Min et al. 2016) both used CNN-based architectures on chromatin accessibility data to predict enhancers.

Here we introduce MuStARD (<u>M</u>achine-learning <u>S</u>ystem for <u>A</u>utomated <u>R</u>NA <u>D</u>iscovery), a flexible Deep Learning framework that can be applied to any biological problem that involves deconvolution of patterns embedded in DNA/RNA sequences. The framework’s flexibility stems from its modular design and minimum input requirements (Figure 1a). The majority of existing algorithms that perform classification tasks in various fields of biological research, pre-miRNA detection for example, rely on extraction of arbitrary features from raw input data. This process often requires significant expertise on the relevant field, it can cause increased computational overhead and most importantly it frequently introduces biases that can severely affect the training of robust models. MuStARD is a feature-agnostic DL framework that utilizes convolutional layers to scan the input data avoiding manual feature extraction. MuStARD is able to work on three types of input or any combination of those, raw DNA sequence, RNAfold (Lorenz et al. 2011) derived secondary structure and PhyloP (Pollard et al. 2010) basewise evolutionary conservation score of the corresponding sequence.

**Figure 1:**
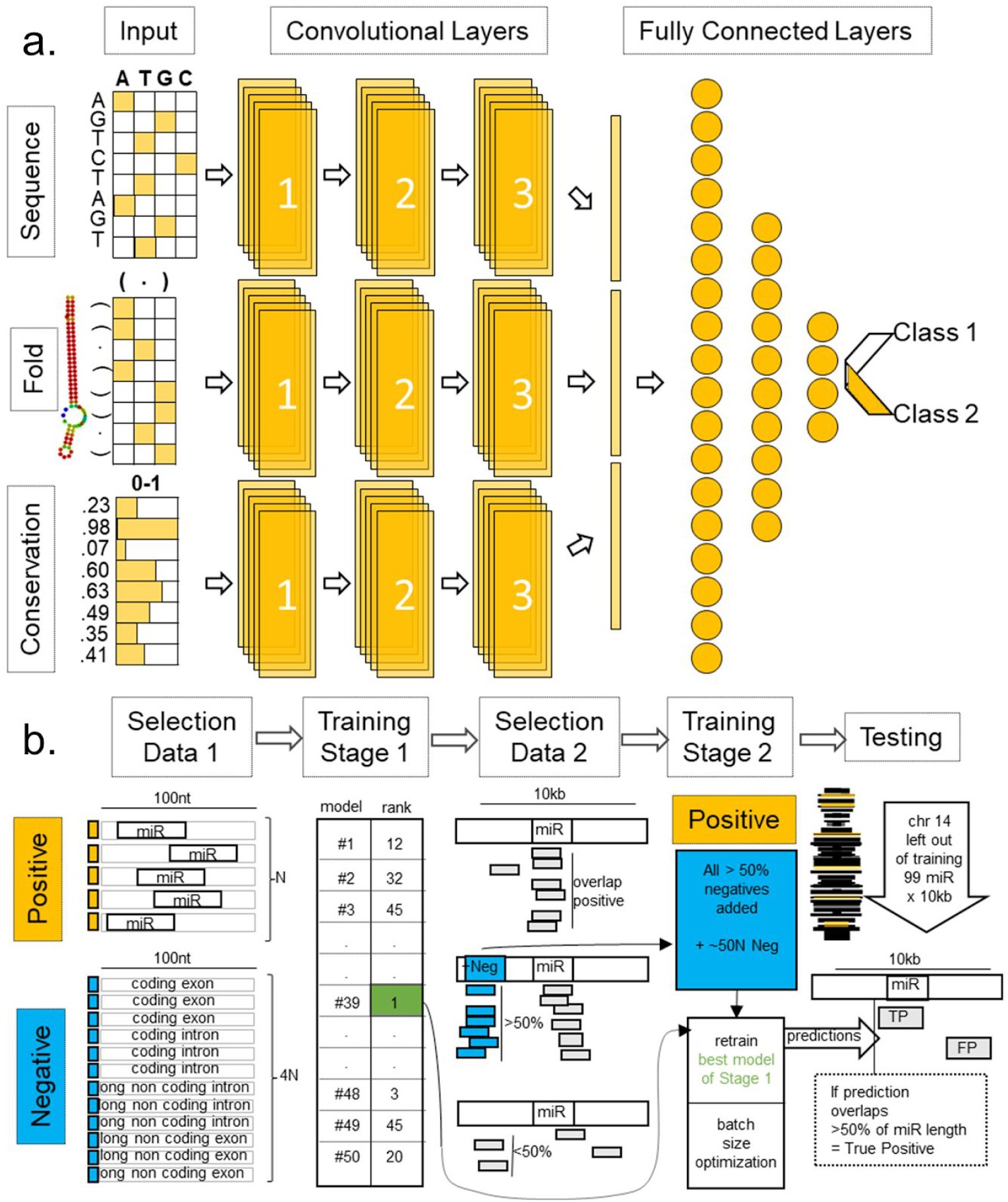
Overview of MuStARD modular architecture and iterative training pipeline. a) MuStARD is able to handle any combination of either raw DNA sequences, RNAfold derived secondary structure and basewise evolutionary conservation from PhyloP. DNA sequences and RNAfold output are one-hot encoded while PhyloP score is not pre-processed. Each feature category is forwarded to a separate ‘branch’ that consists of three convolutional layers. The computations from all branches are concatenated prior to being forwarded to the fully connected part of the network. b) The training pipeline of MuStARD consists of two steps. Initially, pre-miRNA sequences are randomly shuffled to exonic and intronic (protein-coding and lincRNA genes) regions of the genome to extract equal sized negative sequences with 1:4 positive to negative ratio. This process is repeated 50 times to facilitate the training of equal number of models. The performance of each model is assessed based on the test set and all false positives that are supported by at least 25 models are extracted. This set of false positives is added to the negative pool of the best performing model to create an enhanced training set. The enhanced set is finally used to train the final MuStARD model.

## RESULTS

### Evaluation of Input Data Combinations on pre-miRNA prediction

In order to evaluate the importance of the three input data branches on predictive value, we trained MuStARD models using all combinations of one, two and three inputs (sequence (S), conservation (C), and folding/secondary structure (F)) using weighted classes. An additional model was trained for the combination of all three inputs without class weights by disabling the Keras class weights option (MuStARD-mirSFC-U). Each of these models underwent independent hyperparameter optimization for optimal batch size (Supplementary Table 1).

We compared the performance of MuStARD on all combinations of input data for the pre-miRNA prediction dataset. As expected, scanning test sequences with various models shows that models including a higher number of meaningful input data branches perform better in retrieval of pre-miRNAs. The model trained on secondary structure and conservation was the best performing two-input model. This result aligns with the identification of pre-miRNA hairpins by the Microprocessor complex during the biogenesis of miRNAs primarily by characteristics of their secondary structure rather than sequence (Roden et al. 2017) and the fact that pre-miRNAs have highly conserved regions corresponding to the mature miRNA sequences. Surprisingly, the non-balanced model (MuStARD-mirSFC-U) performs best out of all model combinations including the balanced three input model (All comparisons Suppl. Figure 1). Since MuStARD-mirSFC-U outperforms all other models in retrieval of pre-miRNAs, we will only report results for this model in the following evaluations.

### Evaluation of pre-miRNA prediction algorithms on chromosome 14 scanning

While training MuStARD models, we left-out the entirety of chromosome 14 as a final benchmarking/validation set that could be fairly used to evaluate MuStARD’s performance against the current state of the art in pre-miRNA prediction algorithms. The question of accurate pre-miRNA prediction has been thoroughly researched since there are currently over 30 published pre-miRNA prediction algorithms indexed in the OMICtools (Henry et al. 2014) repository. The majority of these studies could not be coerced to run on our benchmarking dataset (See Methods for details). We managed to run and evaluate five state of the art programs: HuntMi (Gudyś et al. 2013), microPred (Batuwita and Palade 2009), miPred (Jiang et al. 2007), miRBoost (Tran et al. 2015) and triplet-SVM (Xue et al. 2005). Of these five, only triplet-SVM, miPred and miRBoost provide probabilities as output scores allowing assessment of their performance on multiple score thresholds. HuntMi and microPred provide fixed output score/labels limiting their performance comparison on a fixed threshold (Supplementary Table 2). After evaluating all five algorithms on the validation set, we identified MiPred as the overall optimally performing state-of-the-art algorithm (Suppl. Fig.2), thus for the sake of brevity we will only report direct in depth comparison to MiPred.

Both MuStARD and MiPred report predictions with probability scores, and both programs would as default be used at a score threshold of 0.5. However, at that threshold, MiPred produces an inordinate amount of false positives (Suppl. Fig. 2). For fairness of comparison of program precision, we have set a threshold on Prediction Sensitivity at the point where each program predicts 50% of real pre-miRNAs (Figure 2a). MuStARD exhibits consistently higher Precision for any level of Sensitivity (Figure 2b,c) and at a strict threshold where 33% of real pre-miRNAs can be annotated it produces on average one False Positive prediction per 800,000 scanned nucleotides (Figure 3d) outperforming MiPred by an order of magnitude.

**Figure 2:**
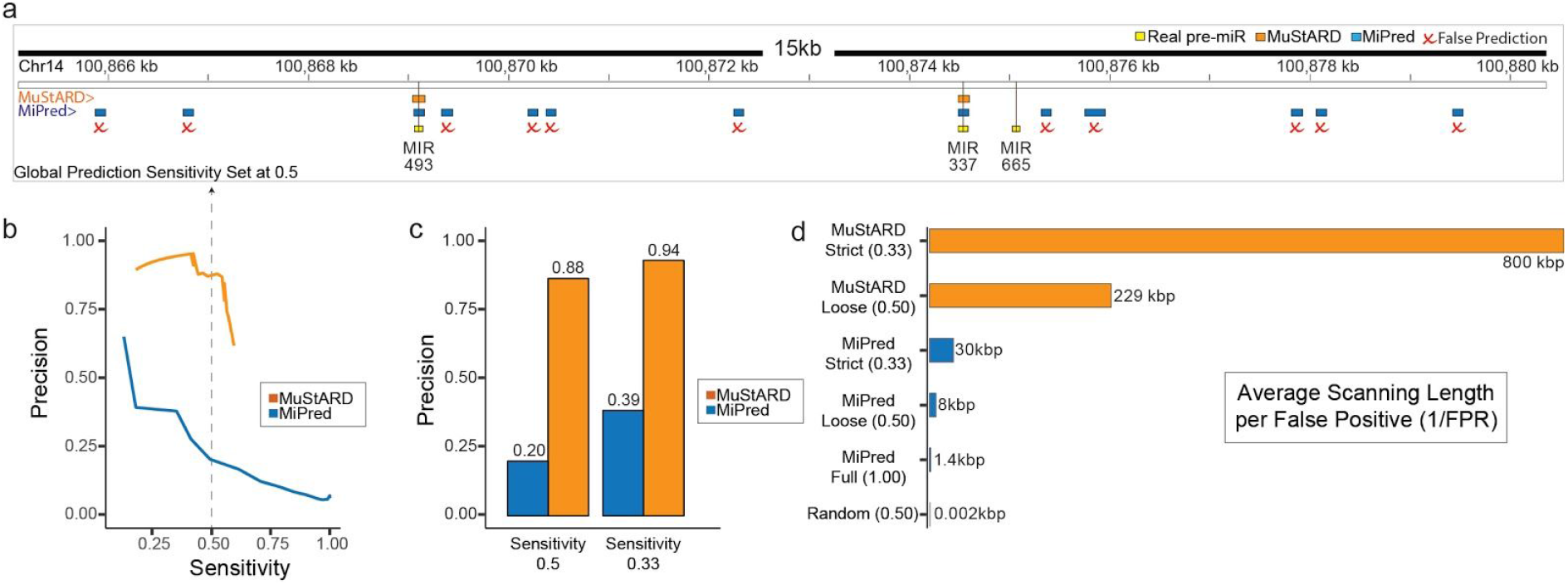
Evaluation of MuStARD human predictions against state of the art prediction algorithm. a) Genome browser visualization of each algorithm’s performance on the scanning windows in a 15kb locus hosting three pre-miRNAs on the left-out chromosome 14. Both evaluated programs have been benchmarked at scores that give Sensitivity of 0.5 over the left-out chromosome. MuStARD correctly predicts 2/3 of the annotated pre-miRNAs, same as MiPred. MuStARD produces no False Positive predictions, compared with 11 of MiPred (marked with red x). b) Precision-Sensitivity curve of MuStARD and MiPred over scanned areas of the left-out chromosome 14. c) Precision of MuStARD and MiPred at loose (sensitivity 0.5) and strict (sensitivity 0.33) thresholds. d) Average length in thousands of base pairs for finding each False Positive prediction on the left-out chromosome. Showing MuStARD at Strict and Loose thresholds, and MiPred at Strict, Loose, and Full (score 0.5 - sensitivity ~1) thresholds, and Random prediction (threshold sensitivity 0.5).

**Figure 3.**
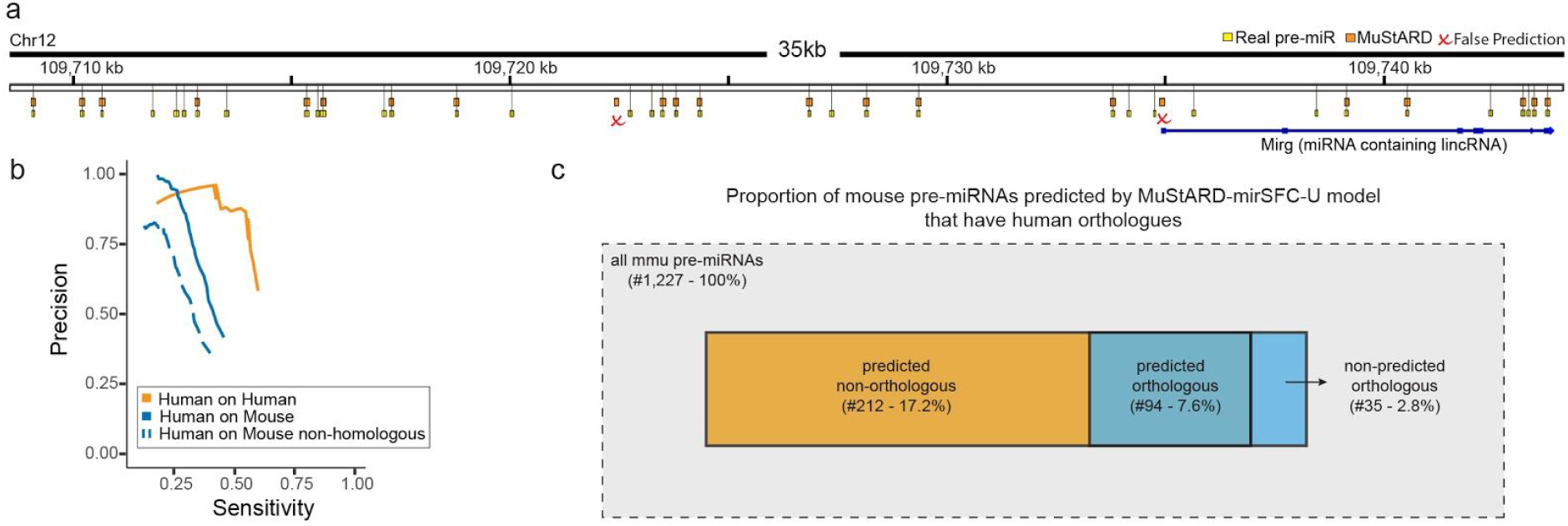
Prediction of Mouse pre-miRNAs by Human trained model. a) Genome browser visualization of MuStARD performance on the scanning of a 35kb locus hosting 36 pre-miRNAs. MuStARD correctly identifies 20/36 pre-miRNAs with 2 False Positives, out of which one falls on the first “exon” of a long non coding RNA Mirg annotated as “miRNA containing lincRNA”. b) Precision-Sensitivity curve of Human trained MuStARD predictions on Mouse pre-miRNAs. Orange line shows the model prediction on Human for reference (same as Figure 2b). Solid blue line shows the prediction on all Mouse pre-miRNAs, and dashed blue line shows the prediction on Mouse pre-miRNAs without a direct human homologue. c) A visualization of the Mouse pre-miRNA evaluation set denoting the number of predicted and non-predicted, orthologous and non-orthologous pre-miRNAs.

### Evaluation of pre-miRNA prediction algorithms on labelled data

The process of genome-wide scanning for pre-miRNAs requires windows of fixed size, a property that perfectly fits the input requirements of Deep Learning algorithms. In reality, the pre-miRNA dataset consists of positive and negative sequences of variable length. These sequences are extended to 100bp prior to MuStARD processing (see Methods for details). However, the majority of existing algorithms instead perform feature extraction and normalization to account for differences on sequence sizes. Using a benchmark dataset of fixed sized sequences should not introduce any biases to comparing the performance of MuStARD and existing algorithms. Nevertheless, we performed an additional comparison based on benchmark sequences (chromosome 14) of the enhanced pre-miRNA dataset without reinforcement. Only for MuStARD, but not for existing algorithms, we applied the extension procedure of these sequences to 100bp. MuStARD models outperform every algorithm in terms of precision. MuStARD-mirSFC-U in particular exhibits unprecedented levels of precision even for score thresholds as low as 0.1 (Supplementary Figure 2 and Supplementary Table 3).

### Inter-species prediction

Having established a substantial increase in precision for intra-species pre-miRNA prediction we evaluated our model on an inter-species prediction. Briefly, we used the best performing pre-miRNA identification model trained on human data, to scan swathes of the mouse genome (in total ~4Mbps) containing 1,227 annotated mouse pre-miRNAs. The inter-species prediction correctly identified pre-miRNAs with a small number of false positives (Figure 3a shows a browser snapshot of a pre-miRNA cluster locus). As expected, the precision of the inter-species prediction was lower than the intra-species validation set (Figure 3b), and even lower for pre-miRNAs that do not have a human homologue as they have lower levels of conservation which is one of our model’s input branches (Figure 3c). MuStARD exhibits exceptional levels of generalisation capacity (Supplementary Table 4) identifying correctly a large majority (94/129) of homologous pre-miRNAs and more than double (212) non-homologous pre-miRNAs.

### snoRNA prediction

Pre-microRNAs are a well defined and well studied class of small non-coding RNAs. Their characteristic secondary structure and conservation pattern made training MuStARD on that dataset a relatively easy task. Despite its performance, MuStARD was not specifically developed for pre-miRNA hairpin detection. Our intention is to provide a highly flexible computational framework that can be applied to the identification of a variety of biological patterns.

To highlight the flexibility of MuStARD, and to understand how it would fare on a more diverse set of patterns, we applied the same training pipeline to train a model of human snoRNA sequences. SnoRNAs are a class of small RNAs with widely varying structure, sequence, and conservation patterns. We experimentally trained a model on all snoRNAs as one class (Figure 4a - orange line) and also for two of the most populous sub-families of snoRNAs, the H/ACA and C/D box families. For the two sub-families we were also able to benchmark against a state-of-the-art snoRNA prediction software developed specifically to identify each of these two categories against background (Figure 4b, Supplementary Table 5). Further, we tested the inter-species capabilities of the MuStARD model, by applying the human-trained snoRNA model to the mouse genome (Figure 4a - blue line).

**Figure 4.**
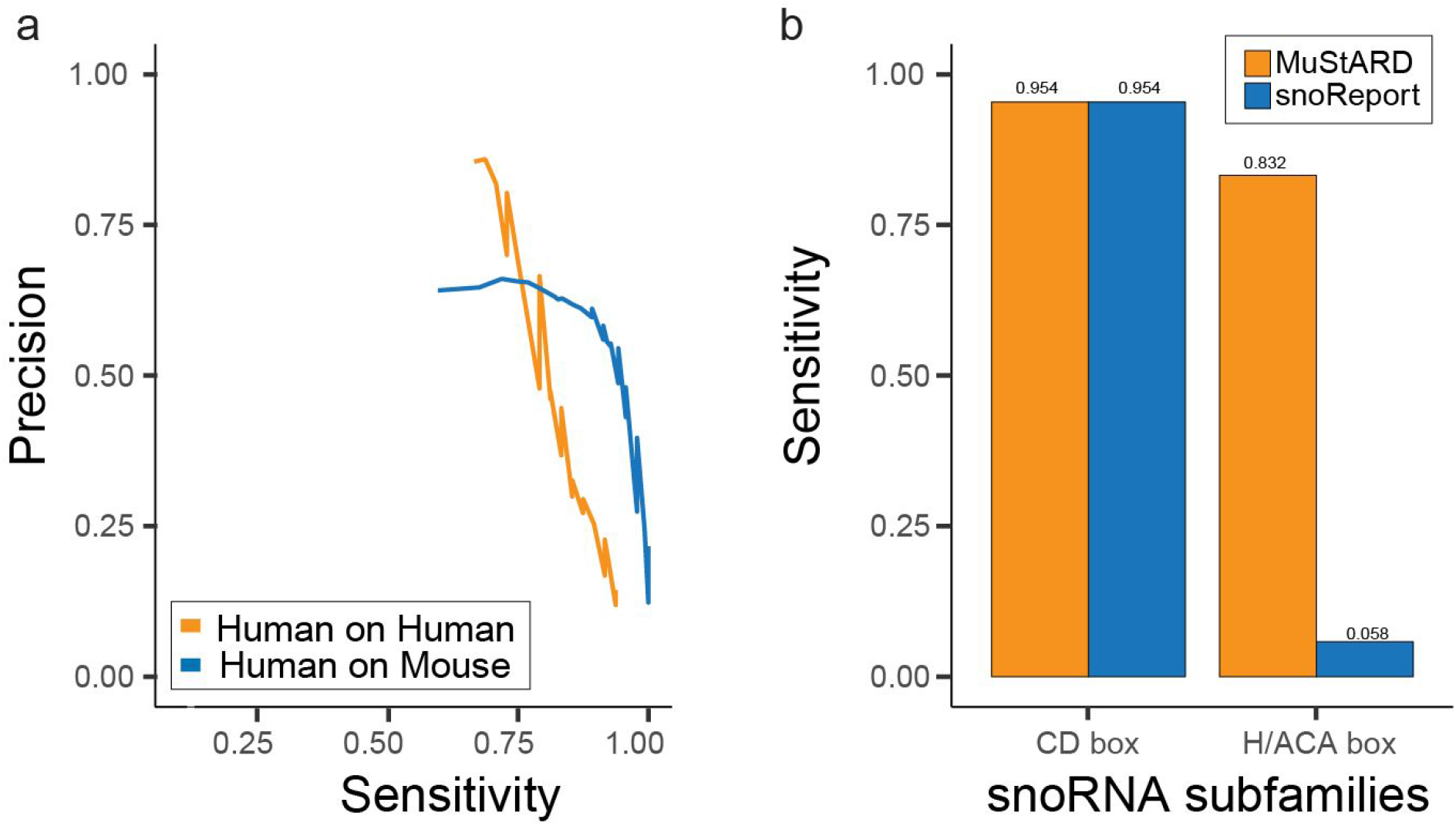
Prediction of snoRNA of CD-box and H/ACA-box subfamilies. a. Precision-sensitivity curves for snoRNA identification of both families. Orange line shows the performance of MuStARD human trained models on left-out human chromosome snoRNAs, blue line shows the performance of the same model on Mouse snoRNAs. b. Comparison of Sensitivity per subfamily against state of the art program snoReport (Precision set at 0.5).

## DISCUSSION

Here we have presented MuStARD, a flexible Deep Learning framework that can be used to identify small RNA genes by example, based on characteristics of their raw sequence, secondary structure and evolutionary conservation, without expertly curated features. We have demonstrated that MuStARD has higher precision over all other methods developed for one specific classification task, and have furthermore, for the first time, successfully attempted a genomic scan in the scale of several million nucleotides. Finally, we point out the potential of MuStARD to annotate classes cross-species (human to mouse) along with moderate evolutionary distances.

An innovative aspect of our method involves the iterative selection of negative examples based on high scoring false positives. Machine Learning methods are only as good as their training set and features. While Deep Learning eliminates the need for expert curated features - some of the pre-miRNA prediction methods utilized up to 700 features (Saçar Demirci, Baumbach, and Allmer 2017) - the need for negative training sets that effectively capture most of the background variation is still needed. We initially prototyped our method with a small set of negatives, four for each real training example. We quickly realized that while our method could separate between these categories easily, it still produced a large amount of false positives in the more realistic scanning test. Training fifty models on fifty sets of negatives improved the performance, but we noticed that specific regions were identified as false positives by a large number of models, i.e. the false positives were not randomly distributed in the background. By enriching our background set with these false positives and retraining the best models in this iterative fashion, we achieved a great leap in performance. An important point for the iterative background enrichment step is that it is fully automated within our method. This allows the method to generalize more easily, since the best background mixture for each class of non-coding RNAs will not be known in advance.

The evaluation of different input modes by itself gave us interesting insight in line with the scientific knowledge of pre-miRNAs. We managed a qualitative ranking of the contribution for each input branch to the final predictive model. Deeper interpretation of the model is beyond the scope of this paper, but is an exciting further field of research. One interesting observation coming from our training, is that class weight balanced training seems to be inferior in accurately predicting pre-miRNAs compared to unbalanced training. Class weight balancing is used in training Deep Learning models so that the model does not attempt to learn the characteristics of a disproportionately populous class while ignoring sparser classes. However, in our more realistic scanning test, one positive example corresponds to at least one hundred negative examples. Our training data with a maximum ratio of approximately one positive example to fifty negatives, although heavily unbalanced is less unbalanced than the realistic testing data. Exploring the class balancing issue will be necessary for the further improvement of the field towards the ultimate goal of genome wide scan prediction.

Using a number of pre-miRNA prediction algorithms for region scanning was time consuming and arduous labor. To calculate hundreds of features on regions spanning less than one percent of the human genome, all other algorithms (with miRBoost being the sole exception) required to group the scanning region into smaller batches of 2000 sequences in order to parallelize the analysis into a computer cluster (MetaCentrum-CERIT). Even so, the computing time for each single batch was approximately 4 days. In contrast, our algorithms was able to scan the mouse benchmark dataset that includes several million base pairs in a few hours on one CPU. With GPU access enabled this process can be even faster. MuStARD has made the possibility of a full mammalian genome scan feasible on a high-end personal computer. However, even with our improved prediction accuracy, the number of false positives identified on a full genome scan would still be disproportionate to the true positives. We will continue exploring improvements and iterative training modification with the goal to achieve genome wide scan capabilities.

Given the increasing number of sequenced genomes becoming available, annotation is lagging. We have demonstrated that MuStARD can be efficiently trained on one species and then used to predict members of the same functional class in another. As a proof of concept we trained models on human pre-miRNAs and snoRNAs and then identified their counterparts in mouse. These species are both well annotated, but have a considerable evolutionary distance. The pre-miRNAs we correctly identified on the mouse genome were enriched in evolutionary conserved pre-miRNAs in human (approximately 30% of our true positive predictions vs 10% of all mouse miRNAs). That said, the majority (70%) of our predicted pre-miRNAs are not homologous to human pre-miRNAs and would not be easily identified by a simple homology search.

We chose pre-miRNAs as a first example because they have well established annotation, consistent secondary structure, conservation and other sequence characteristics. Our second use example was snoRNAs where most of these assumptions fail. The snoRNA class consists of several families that do not share common secondary structure, motif sequences, or conservation profiles. Conservation at large is much less pronounced in snoRNAs compared to pre-miRNAs. Additionally, the size distribution of snoRNAs (118.8bp mean, 59.1bp standard deviation) is much wider than miRNAs (81.9bp mean, 16.9bp standard deviation) making it harder for our method to accurately identify them. Despite these drawbacks, we manage to identify snoRNAs accurately within the human genome and in cross-species scan. In fact, surprisingly, our method seems to identify cross-species validation examples more accurately than left-out human examples. This could possibly be the direct result of having cross-species snoRNAs with conserved sequences to human snoRNAs in the training set. We have opted not to remove these snoRNAs from the cross-species comparison so as to estimate a precision/sensitivity rate closer to what a real scan would produce.

We have developed MuStARD as an easy to deploy, versatile, and extensible method that can run with minimal user input. To make it modular and versatile we used a Keras architecture that can be easily extended from more experienced users. Keras is widely accepted by the Deep Learning community and offers ease of use and several layers of abstraction in terms of code sharing when compared to tensorflow or other lower end frameworks. Extending the architecture with more diverse branches is straightforward. We have already developed templates for reading sequence and secondary structure (one hot encoding) as well as conservation (scores). Adding different types of signal is as easy as downloading a relevant track from UCSC Genome Tracks and asking MuStARD to include it in the training. For basic users, the input has been kept minimal, requiring just a bed file of regions in the functional class of interest, a sequence file of the genome, and a conservation track of the same size. Given these inputs, MuStARD preprocesses the regions of interest, extracts sequences, simulates folding, picks random genomic sequences for background, optimizes hyperparameters and so on until the final model is trained.

## MATERIALS AND METHODS

### Dataset collection and preprocessing

Human (GRCh38) and mouse (GRCm38) genomes and corresponding gene annotations were downloaded from Ensembl v93 repository(Zerbino et al. 2018). Human gene annotation was filtered to only include genes exhibiting a protein-coding or lincRNA biotype. The exons of protein-coding and lincRNA genes were combined to produce two disjoint data sets; parts of the genome that correspond to exons and loci that are represented by introns. Additionally, two separate collections were created by selecting regions marked with the snoRNA biotype. Human and mouse pre-miRNAs were downloaded from miRBase v22 (Kozomara, Birgaoanu, and Griffiths-Jones 2018). Human pre-miRNAs were subsequently filtered based on the experimentally validated information provided in miRBase to keep only high-quality sequences for training. Basewise conservation scores, based on phyloP algorithm, of 99 and 59 vertebrate genomes with human and mouse respectively were downloaded from UCSC genome repository (Karolchik et al. 2004).

### MuStARD training module architecture

The aim of MuStARD is to provide a highly flexible, feature-agnostic computational framework that can be applied in a plethora of Biological problems providing state-of-the-art performance while at the same time having minimal input requirements. To this end, MuStARD has been specifically designed to follow a modular architecture where each module carries out different functionalities that can be run and assessed independently and/or in parallel (Figure 1). The framework is implemented using python for the deep learning aspect, R for the majority of meta-analyses and plotting, and perl for general purpose file filtering, formatting and module connectivity. Users only need to provide bed formatted files as input and the appropriate genome assembly files as well as the wiggle formatted PhyloP evolutionary conservation score files derived from UCSC repository.

The training module of MuStARD is composed of a convolutional architecture based on tensorflow and the Keras functional API. More advanced users can directly add or remove parts of the architecture according to the problem at hand. For the purposes of this study, the chosen architecture consists of 3 convolutional branches that can be dynamically added, removed and combined in multiple ways according to the properties of the corresponding use-case. These branches depict distinct ‘agents’ that are able to independently model different input modes such as raw DNA sequence, RNA secondary structure and evolutionary conservation. Subsequently, the outputs of the convolutional branches are flattened, concatenated and forwarded to the dense part of the architecture that produces the final prediction scores. In every layer output, dropout and batch normalization regularization techniques are applied to improve the generalisation capacity of the network.

Regardless of the chosen network architecture, hyperparameters are known to be notoriously hard to optimize and depending on the complexity of the input, small changes in the hyperparameter selection can greatly affect the results. The training module has been designed to incorporate a grid-search type of approach for finding the optimal combination of hyperparameters. We have chosen to apply grid-search over 4 hyperparameters, the ones that based on our experience are able to greatly affect the results; batch size, learning rate, dropout rate and number of filters in the convolutional layers. Users can freely remove or add hyperparameters into the grid-search process and most importantly adjust the network architecture according to their needs. Each model trained over a different combination of hyperparameters is saved in a separate directory alongside train/validation accuracy/loss plots and a detailed log of the performance in each epoch. This allows users to find the exact combination of hyperparameters that produces the optimal training.

Unless stated otherwise, in all use-cases presented in this study, each convolutional branch consists of 3 convolutional layers. The first convolutional layer in the raw DNA sequence processing branch uses a filter size of 16 nucleotides with stride 1 and no padding, the second layer uses a filter size of 12 and the third layer a filter size of 8. The first convolutional layer in the RNAfold processing branch uses a filter size of 30 nucleotides with stride 1 and no padding, the second layer uses a filter size of 20 and the third layer a filter size of 10. The first convolutional layer in the evolutionary conservation processing branch uses a filter size of 20 nucleotides with stride 1 and no padding, the second layer uses a filter size of 15 and the third layer a filter size of 10. The outputs of the convolutional branches are flattened and concatenated before being forwarded to the dense part of the network that includes 3 layers of 100, 75 and 50 nodes respectively. All layers use leaky ReLu activation except the final prediction layer that uses the softmax function. The chosen optimizer is SGD with Nesterov momentum set at 0.9. All models were trained over 600 epochs after enabling early stopping with patience set at 40 and delta at 0 with a learning rate of 10^−4^.

### MuStARD prediction module

The prediction module of MuStARD framework has been explicitly designed to facilitate both long region scanning and static assessment of specific loci. In the case of long region scanning, users are able to select the appropriate parameters such as the window size (it should match with the training window size), sliding step and the model that will be used for scoring each window. The framework includes standalone code for generating bedGraph tracks that can facilitate the visualization of results in any genome browser as well as code for creating ‘hotspots’ of positive predictions and for evaluating the results based on custom tracks and/or annotations. In the case of static assessment of specific loci, the prediction module provides a bed formatted file that included the score of each region in the 5th column.

### Learning process of pre-miRNA and snoRNA detection MuStARD models

As described in previous sections, the training (Figure 1) of the pre-miRNA recognition model was based on experimentally verified human precursor sequences from miRBase. Only pre-miRNAs with size less than 100bp were used to form the positive set resulting in 579 sequences that were separated into the necessary training (484 hairpins corresponding to all chromosomes except 2, 3 and 14) and validation (51 hairpins corresponding to chromosomes 2 and 3) sets required by Deep Learning frameworks for the training process. 99 hairpins from chromosome 14 (no experimental validation evidence or size distinction) were completely left out from the training procedure and were utilized for testing the performance of pre-miRNA detection algorithms. The negative set was formulated with bedtools 2.27.0v(Quinlan and Hall 2010) ‘shuffle’ mode using the positive set on the exon/intron genomic segments described in previous section. For each positive instance 4 equally sized negatives were randomly selected from protein- and non-coding exonic as well as intronic regions, 1 for every category. This process was repeated 50 times in total creating 50 different training/validation sets that were used to train an equal amount of distinct preliminary models. Hyperparameters were fixed at 256 batch size, 0.2 dropout rate, 0.0001 learning rate and 80/40/20 number of filters in the 3 convolutional layers of each branch and the class weight option in Keras was enabled. Based on this repetitive negative shuffling configuration we ensured that a reasonable balance between training time as well approximating sequence and evolutionary conservation variation in background or non-precursor genomic loci was maintained. One of our objectives was to optimize the genome scanning process. The majority of existing algorithms utilize positive sequences that are fixated around the center of pre-miRNAs. However, in genome scanning scenarios there will always be instances in which part or the whole hairpin sequence will not be located in the center of the scanning window. This phenomenon might heavily affect the secondary structure of the RNA sequence corresponding to each window and therefore the generalisation capacity of the model. To overcome this problem, the MuStARD training module has been equipped with an optional ‘reinforcement’ feature that generates copies of the input instances with randomly placed positive or negative sequences within the 100nt sequence. For the purposes of this study, the number of reinforced instances for every input sequence during the pre-miRNA training process was 5.

Ideally, if the combination of using intronic/exonic regions as a background sequence pool and the 1:4 positive to negative ratio was enough to fully capture the non-precursor sequence variation in the 3 input feature space (raw DNA sequence, secondary structure and basewise evolutionary conservation) then a near perfect performance in terms of both precision and sensitivity would be achieved in a scenario where all 50 preliminary models are used to scan the genome for predicting pre-miRNA sequences. To test this hypothesis all human pre-miRNAs were extended by +/− 5,000bp and the resulting regions were merged in the case of strand specific overlaps. Both strands of the merged loci were scanned with all 50 preliminary models using a window of 100bp and a stride of 10bp. This resulted in a benchmark dataset of 33.2 million bp divided into 3.2 million overlapping 100bp windows. For each model, out of the 3.2 million windows only those exhibiting a score above 0.5 were retained to form ‘hotspots’ of positively predicted regions after merging cases of strand specific overlaps. These regions were subsequently cross checked with the annotated pre-miRNAs to extract performance metrics in the 0.5-0.9 score range for every preliminary model.

False positive predictions based on a 0.5 score threshold were kept only if they were supported by 25 out of 50 preliminary models and did not overlap with any negative instance used to train these models. The resulting 23,750 false positive loci were added to the negative dataset of the best performing preliminary model. These false positives represent regions of the genome that were not captured by the process of ‘shuffling’ positive instances to exonic/intronic loci and exhibit feature characteristics that are more similar to positive than negative instances. This process assisted in establishing an enhanced set of sequences that was used to train the final pre-miRNA detection model that was selected through performing a hyperparameter space grid-search over the batch size and the Keras option of training with/without class weights (Supplementary Table 1). The class weights option in Keras enables the equal contribution of all classes during the training of unbalanced datasets. The remaining hyperparameters were not changed.

This process was repeated 6 times to train, with the Keras class weights option enabled, an equal number of distinct MuStARD pre-miRNA detection models composed of different input combinations; raw sequence with secondary structure and conservation (MuStARD-mirSFC model), raw sequence and conservation (MuStARD-mirSC), raw sequence and secondary structure (MuStARD-mirSF), secondary structure and conservation (MuStARD-mirFC), secondary structure only (MuStARD-mirF) and sequence only (MuStARD-mirS). For the combination of raw sequence, secondary structure and conservation, we have trained an additional model after disabling the class weights option in Keras (MuStARD-mirSFC-U model). For MuStARD-mirSFC model, the optimal (balance between precision and sensitivity) batch size was 1024, 256 for MuStARD-mirSFC-U, 1024 for MuStARD-mirSC, 256 for MuStARD-mirSF and MuStARD-mirFC, 512 for MuStARD-mirF and MuStARD-mirS. The procedure for evaluating the performance of each model is described in the following section.

For the purposes of the second use-case presented in this study, the same pipeline was used to create three MuStARD snoRNA detection models, one for detecting the CD box snoRNA subspecies (MuStARD-snoSFC-U-CDbox), one for H/ACA box (MuStARD-snoSFC-U-HACAbox) and one for detecting both subspecies at the same time (MuStARD-snoSFC-U). For training the MuStARD-snoSFC-U-CDbox model we removed CD box snoRNAs with size more than 130nt. For the validation set we used 28 sequences originating from chromosomes 3, 4, 5, 20, 21 and 22. For the training set we utilized 208 sequences located in the remaining chromosomes except 22 snoRNAs from chromosomes 1 and 2 (no size distinction for the creation of this set) that were used for testing the performance of snoRNA prediction algorithms. For training the MuStARD-snoSFC-U-HACAbox model we removed H/ACA box snoRNAs with size more than 150nt. For the validation set we used 29 snoRNAs originating from chromosomes 2, 3, 4, 20, 21 and 22. For the training set we utilized 94 sequences located in the remaining chromosomes except 17 snoRNAs from chromosome 1 (no size distinction) that were kept for testing. For training the MuStARD-snoSFC-U model we removed H/ACA and CD box snoRNAs with size more than 150nt. For the validation set we used 42 snoRNAs originating from chromosomes 3, 4, 20, 21 and 22. For the training set we utilized 317 sequences located in the remaining chromosomes except 48 snoRNAs from chromosomes 1 and 2 (no size distinction) that consisted the testing set.

### Testing on genomic region scanning data

The process of testing algorithms on a static labelled dataset can often provide misleading results about performance especially in cases of models that have been designed for genome-wide scanning. Such ‘stress’ tests are often also able to unveil interesting aspects about the computational complexity and the time required by algorithms to complete a task. To this end, 99 human pre-miRNAs (no size or quality distinction) located on chromosome 14 were extended by +/− 5,000bp and the resulting regions were merged in the case of strand specific overlaps. Both strands of the merged loci were scanned with all MuStARD’s final pre-miRNA detection models (Supplementary Table 1) as well as with existing algorithms (Figure 2, Supplementary Table 2) using a window of 100bp and a stride of 5bp. This resulted in a scanning benchmark dataset of 1 million bp divided into 208,708 overlapping 100bp windows.

Two distinct strategies were employed to assess the performance of each algorithm. In the first approach, each window was assessed independently (method A) while in the second only windows exhibiting a score above 0.5 were retained to form ‘hotspots’ of positively predicted regions after merging cases of strand specific overlaps (method B) or overlaps regardless of strand (method C). These regions were subsequently cross checked with the 99 annotated chromosome 14 pre-miRNAs to extract performance metrics in the 0.5-0.9 score range, when possible. In all scenarios, positive predictions were considered true positives (TPs) if they covered at least 50% of the overlapping annotated pre-miRNA’s size. HuntMi(Gudyś et al. 2013) and microPred(Batuwita and Palade 2009) algorithms only provide hard labelled results instead of a probabilistic score, therefore they were not included in the graphs. However, to facilitate a fair comparison between all algorithms, performance metrics based on all methods (A, B and C) were extracted at a fixed score threshold of 0.5 (Supplementary Tables 2, 3 and 4).

For assessing the performance of MuStARD-snoSFC-U, MuStARD-snoSFC-U-CDbox, MuStARD-snoSFC-U-HACAbox, snoReport-CDbox and snoReport-HACAbox models human snoRNAs on the test chromosomes (see in previous section) were extended by +/− 10,000bp and the resulting regions were merged in the case of strand specific overlaps. Both strands of the merged loci were scanned with different window sizes (depending on the model) and a stride of 5bp. For comparing the snoRNA detection models, only method C was applied.

Method C was also applied to annotated mouse pre-miRNAs and snoRNAs using MuStARD’s SFC-U models and snoReport in the case of snoRNAs (Figures 3 and 4, Supplementary Tables 4 and 5).

### Testing on labelled data

For the purposes of testing the final pre-miRNA detection model on labelled data and comparing with existing algorithms, all positive and negative instances located on chromosome 14 were used from the enhanced data set described in the previous section after removing sequences with size less than 100bp. The total number of positive instances in the test set was 44 and the total number of negatives 893 (Supplementary Figure X, Supplementary Table 3).

### Application of existing algorithms

There are over 30 pre-miRNA prediction algorithms listed in OMICtools repository. The majority of these studies provide access to the trained models only through web-server interfaces which allow a small number of sequences to be processed at once. Only a handful of studies provide stand-alone implementations that can be downloaded and applied on benchmark datasets locally. However, a small fraction of these implementations are able to properly function and provide results.

We only managed to assess the prediction efficiency of HuntMi, microPred, MiPred, triplet-SVM, and MirBoost on our benchmark datasets. HuntMi and microPred tools do not support parallelization, and the average processing time for a sequence of 100nt is 3 minutes. The scanning benchmark sequences were grouped into 100 bins to faster the analysis for HuntMi and microPred. Also, microPred random sequence generation parameter setting was 500. Each bin was analyzed independently by HuntMi and microPred on virtual machines provided by the MetaCentrum-CESNET supercomputer cluster. MiRBoost’s SVM model was re-trained to support probabilistic output using the dataset included in the code repository and parameters ‘svm-train -h 0 -c 8.0 -g 0.125 -w1 1 -w-1 1 -b 1’. Then miRBoost was applied on our benchmark dataset with parameters ‘miRBoost -d 0.25’. For triplet-SVM, we initially applied RNAfold on our benchmark dataset with parameters ‘RNAfold --noPS --noconv --jobs=10’ and the output was forwarded to the triplet-SVM perl script with parameters ‘triplet_svm_classifier.pl 22’ that pre-processes the data and reformats it for the final prediction modules that requires libsvm. The final triplet-SVM results were obtained using svm-predict with parameters ‘svm-predict -b 1’. MiPred was applied on the benchmark dataset with default parameters. For the snoRNA use-case snoReport was used with default settings.

### Assessing MuStARD’s ability to detect non-human-homologous pre-miRNAs in mouse

Mouse hairpins regions of miRNA transcripts (N=1,227) were derived from the miRBase database; orthologous miRNA (N=129) between mouse and human were retrieved from the Ensembl BioMart hub(Kinsella et al. 2011). Initially, accurate MuStARD predictions (true positives) were recognized as overlapping with mouse hairpins regions through bedtools intersect v2.27.1. Subsequently, non-human-homologous pre-miRNAs were distinguished as the negative intersection between accurate MuStARD predictions and the human orthologous miRNA dataset. Bedtools options ‘same strandedness’ and ‘overlaps=0.5’ were used in both cases (-s and -f, respectively).

### Software and hardware requirements

MuStARD is developed in Python 2.7 for the Deep Learning aspect (tensorflow 1.10 and Keras 2.2.2), R for visualizing the performance and Perl for file processing, reformatting and module connectivity. Full list of dependencies can be found on MuStARD’s gitlab page. MuStARD is able to execute either on CPU or GPU depending on the underlying hardware configuration by taking into advantage tensorflow’s flexibility. The framework has been designed to maintain a minimal memory footprint thus allowing the execution even on personal computers. Running time heavily depends on input dimensionality, number of instances in the training set, learning rate and GPU availability.

## AVAILABILITY

MuStARD is an open source Deep Learning framework available in GitLab (<u>gitlab.com/RBP_Bioinformatics/mustard</u>).

## SUPPLEMENTARY DATA

Supplementary Data are available at NAR online.

## ACKNOWLEDGEMENT

We would like to acknowledge ‘MetaCentrum NGI’ for providing computational infrastructure necessary for running parts of the analyses presented in this study.

